# Weddell seals near the fastest melting glacier in Antarctica prefer shallow, coastal and partially ice-covered waters

**DOI:** 10.1101/2022.10.24.513500

**Authors:** Guilherme A. Bortolotto, Karen J. Heywood, Lars Boehme

## Abstract

Antarctic ice-shelves melting is accelerating, and the highest rates for the continent are observed in the Amundsen Sea, where Weddell seals are understudied. By modelling an unrivalled telemetry tracking dataset, we show that this top predator in the area is more likely found in shallower waters, closer to the coast, half-covered by ice and where the temperature at the ocean bottom is around 0.5°C or colder. Their distribution during winter is more restricted to coastal areas in the eastern Amundsen Sea and spread out in warmer months. That fluctuation reflects the drastic variation in their habitat illustrating their potential sensitivity to the upcoming impacts of climate changes. We discuss their overlap with Antarctic toothfish fisheries in the area and the potential consequences of global warming to the ecology of the seals.

## Introduction

The melting of ice shelves in Antarctica has accelerated recently (Paolo, Fricker, and Padman 2015) which can result in a global sea level rise of over 10 meters in a few decades or centuries (DeConto and Pollard 2016). The highest levels of melting for the continent are observed in the Amundsen Sea (Paolo, Fricker, and Padman 2015; Rignot et al. 2019), and it is certain that coastlines around the world will change drastically (Le Cozannet et al. 2019), yet very limited information exists on how the local environment dynamics are being affected (Yoon et al. 2022; Zheng et al. 2021; Dotto et al., n.d.), with little effort on understanding local species ecology.

The iconic Weddell seal (*Leptonychotes weddellii*) is the southernmost breeding mammal in the planet, requiring Antarctica’s fast-ice habitat for reproduction. Despite intense biological and ecological investigation, much of the research effort for the species was in the Ross Sea (Goetz et al. 2017; Rotella et al. 2012), and recent research suggests the presence of the species in the Amundsen Sea to be relatively low when compared to the rest of the continent (LaRue et al. 2021). Mapping of Weddell seal local (Figure 10 in Bengtson et al. 2011) and circumpolar (Figure 1 in (LaRue et al. 2021) distribution even indicate the species to be absent in the eastern limit of the Amundsen Sea. Access to that area for *in situ* observations can be difficult due to persistent floating ice cover (Hogan et al. 2020), a constrain that telemetry tagging of seals can help overcome.

**Figure 1.**
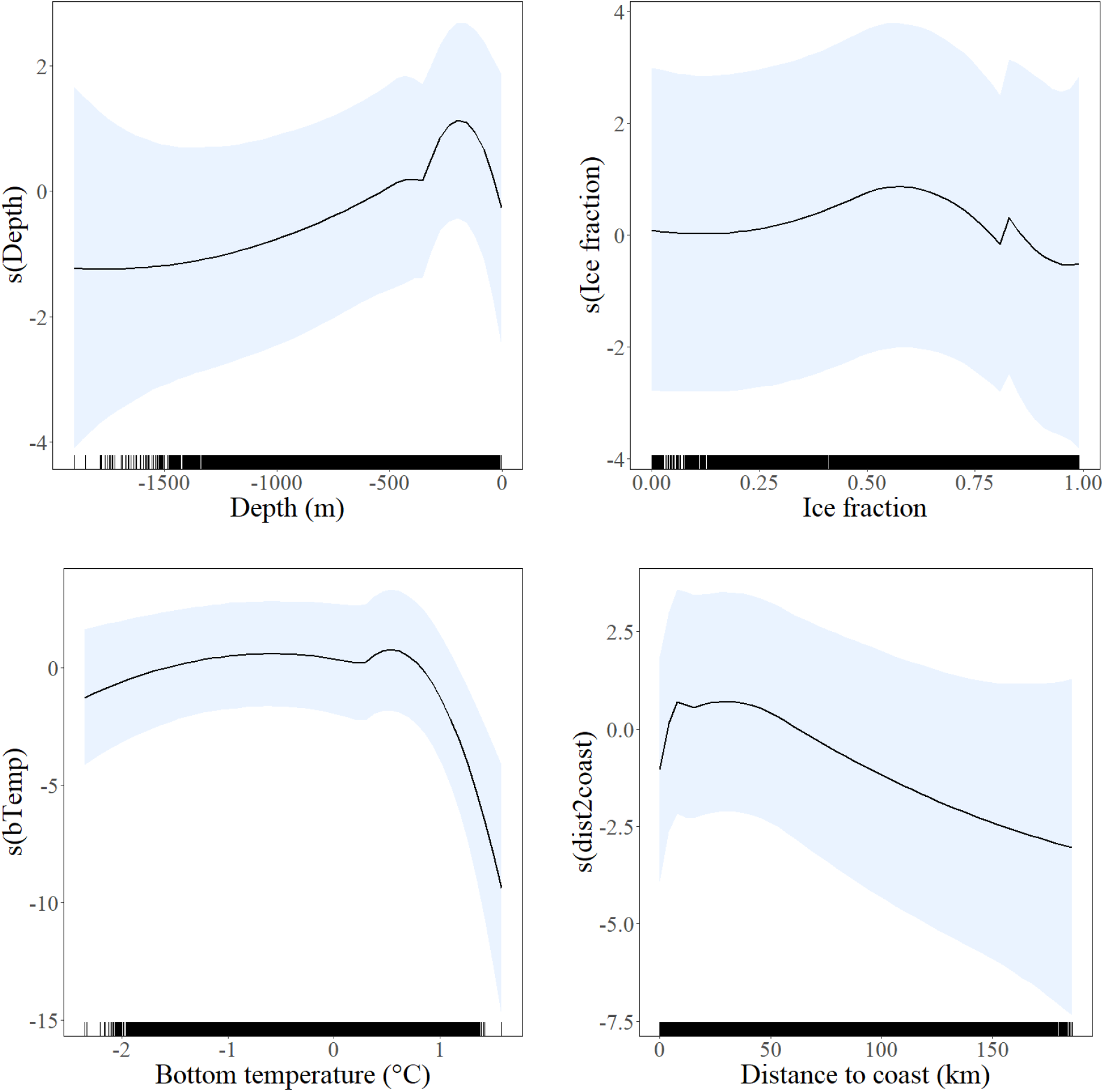
Habitat use model results –. Smooth functions fitted to covariates retained in the selected habitat use model. R software code for plotting smooth functions resulting from GEE model fitting were adapted from (Pirotta et al. 2011).

A major component of the diet of Weddell seals is the Antarctic toothfish (Ainley et al. 2021), a large predator itself, largely target by fisheries in the Amundsen Sea (Queirós et al. 2022). All Antarctic toothfish fisheries are regulated by the Commission for the Conservation of Antarctic Marine Living Resources taking an ecosystem-based and conservative approach (Hanchet et al. 2014; CCAMRL 2022). In that context, information on the ecology of seals that predate on them in an area likely to suffer drastic changes in the next years due to climate change, can further assist informing conservation strategies for multiple species in that region.

To confirm the importance of the Amundsen Sea for Weddell seals, by illustrating their distribution and habitat use, we modelled telemetry tracking data to show how that relates to important environmental covariates in the fast-changing habitat of the Amundsen Sea. Understanding the ecology of this predator in that area will be key to evaluate how environmental changes may affect local species communities, and the potential interaction with fisheries in the Amundsen Sea.

## Results and Discussion

### Tracking data

Tracking information from 26 seals were retained for analysis after data processing and filtering steps, including tracks from 15 males, 8 females and 3 individuals of undetermined sex. There were 8 tracks from seals tagged in 2019, 6 in 2020 and 12 in 2022. After data filtering and pre-processing, 3580 presences and 17900 pseudo-absences were used for modelling the habitat use of Weddell seals (Supplementary Fig. 1). This is a relatively large animal telemetry tracking dataset (Börger et al. 2006), and the first to be analysed for habitat use and distribution for Weddell seals in the Amundsen Sea.

### Habitat use

The selected habitat use model included covariates depth, surface ice fraction, distance to the coast and temperature at the ocean bottom (Figure 1). No interannual temporal effects were identified, with none of the factor covariates (month, year or accessibility polygon) retained in the selected model. Residual serial autocorrelation was high (Supplementary Fig. 4), highlighting the need to use Generalised Estimation Equations for model fitting and adequate consideration of coefficient estimated uncertainties. Model performance was good, with a confusion matrix showing 71.5% of correct predictions (i.e., 75.3% correctly predicted for presences and 70.8% for pseudo-absences) and the area under the ROC curve statistic was 0.8.

Habitat use modelling results partially corroborate previous investigations of habitat driven heterogeneity of Weddell seals distribution: pup-rearing seals were observed to more likely haul out very close (less than 2 km) form the coastline (LaRue et al. 2021), but that did not effectively consider distribution of animals at-seas, while swimming or diving (i.e., foraging). Our results also indicate that Weddell seals are more likely found closer to the coast, however a peak in occurrence is observed at around 50 km from the nearest point inland (Figure 1). Also, the only penguin colony in the area is likely the one found in the islands near Canisteo Peninsula, and because of the likely confounding effect with distance to the coast, we did not consider the influence of the presence of colonies of competitor species (LaRue et al. 2021). However, if the presence of that Adelie penguin colony is somehow related to the presence of seals in that area, is a question for future investigation. Since fast-ice is necessarily near the coast, and we found a Weddell seals to be more likely found in areas where half of the surface was covered by ice (Figure 1), that is consistent with their preference for fast-ice to haul-out (Bengtson et al. 2011), near cracks in the ice (LaRue et al. 2021). Despite the fact that we did not differentiate between the type of ice in the “ice fraction” covariate (e.g., pack ice vs. fast-ice vs. icebergs), understandably Weddell seals prefer areas with both easy access to floating ice, for protection from predators, and to open water, for foraging. We provide here more general insights into habitat use for Weddell seals than previous studies, not focusing on a specify month when “low level of foraging” is expected (the pup-rearing month, November; LaRue et al. 2021).

Relations between the depth (<300m) and the temperature at the ocean bottom (∼0.5°C or colder) and higher probabilities of seal occurrence can likely be explained by the prey availability (LaRue et al. 2021). Increasingly evidence suggest that toothfish is a key prey species to Weddell seals (Ainley et al. 2021), and our results suggest spatiotemporal overlap between out seal distribution and a Small Scale Research Unit (SSRU) managed by Commission for the Conservation of Antarctic (CCAMRL 2022). Most Antarctic tooth fishing in hat SSRU happens during warmer months (Queirós et al. 2022), when we demonstrate seals to be more spread out in the area (Figure 2). Fishing effort and seal distribution information must be jointly considered if to successfully address conservation issues for either species in the area.

**Figure 2.**
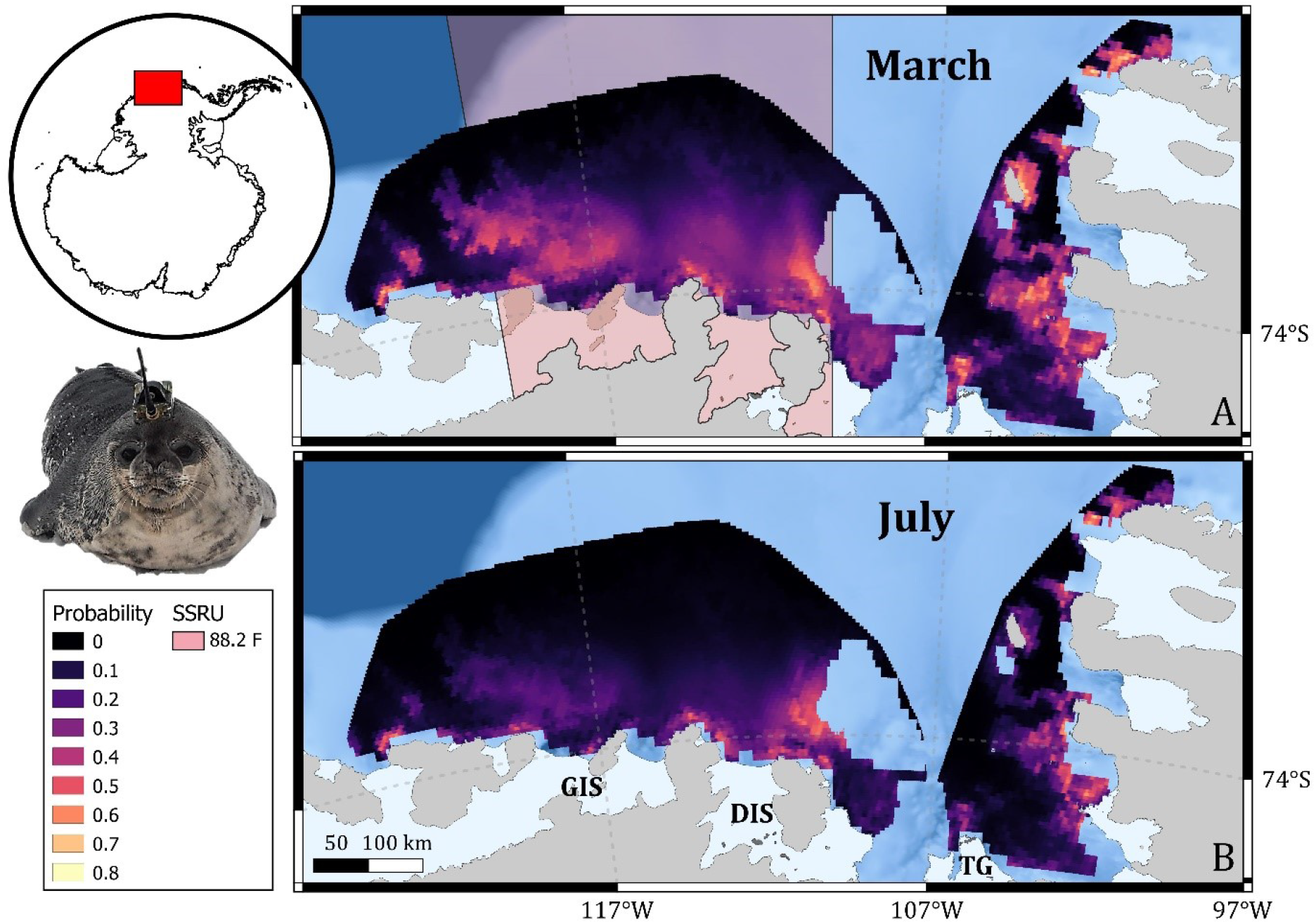
Distribution of Weddell seals – relative probability of occurrence. A) Predictions for March to illustrate distribution in warmer periods (Small Scale Research Unit, SSRU, 88.2F is shown for reference) and B) in July for winter distribution. The probability of seal occurrence was predicted from a habitat use model including water depth, distance to the coast, temperature at ocean bottom and surface ice fraction, with monthly observations averaged for the last 10 years. Predictions for all months can be found in Supplementary Fig. 5.

The probability of seal occurrence was heterogeneous in space and between months (Supplementary Fig. 5) with predictions uncertainty not varying temporally (Supplementary Fig. 6). Seals concentrated directly to the north of the Getz and Dotson ice-shelves and of Martin and Bear peninsulas, and near the coast along the eastern limit of the Amundsen Sea during colder months (Figure 2B). Their distribution spreads out during warmer months (Figure 2B). The only other study to focus on the presence of Weddell seals in an area similar to ours (Bengtson et al. 2011) had visual survey effort limited to March 2000, and suggests an absence of the species in the area, which is clearly not the case.

The post-calculated dosage of Zoletil® averaged 0.5 mg/kg when the intravenous (IV) route was used, and 1 mg/kg when intramuscular (IM). One of the seals that received an initial IV dosage of 0.7 mg/kg (“Akira”, tagged in 2020), and two that received IM initial dosages of 1.1 mg/kg and 1.2 mg/kg (“Manoela” and “Cassandra”, both in 2022), presented apnoea (McMahon et al. 2000) and were successfully mechanically ventilated using an endotracheal tube attached to manual 2-way air pump.

## Methods

Distribution and habitat use of Weddell seals in the Amundsen Sea were investigated through statistical modelling of animal tracking data and environmental covariates.

### Ethics statement

Seal tagging in Antarctica was conducted under permits 29/2018, 03/2019-20 and issued by the Foreign and Common Office of the UK. Project TARSAN has ethical approval from the School of Biology Ethics Committee of the University of St Andrews.

### Data collection

Seals were tagged during research cruises aboard the RVIB Nathaniel B. Palmer, as part of the International Thwaites Glacier Collaboration. Seal tracks were collected using Conductivity-Temperature-Depth Satellite Data Relay Loggers (CTD-SDRLs; Boehme et al. 2009) manufactured by the Sea Mammal Research Unit Instrumentation Group (SMRU-IG), in St Andrews, Scotland, UK. A dataset from 39 Weddell seals (23 males, 13 females and 3 of undetermined sex), tagged in 2019 (n = 12), 2020 (n = 9) and of 2022 (21) were available, representing the largest ever Weddell seal telemetry tracking dataset from the Amundsen Sea.

Seals were captured after their annual moult, in February-March, and chemically restrained using Zoletil® (Bayer®). A conical canvas bag was used for physical restraint, either before an IV (via the extradural intravertebral vein; Mcmahon et al. 2000), or after IM application (via dart to the lumbar muscles) of the tranquilizer. The weight of the seals was estimated by eye for calculating the volume of anaesthetics to be used, following dosage recommended in the literature (i.e., 0.5 mg/kg intravenously, IV, as per Wheatley et al. (2006); 1 mg/kg intramuscular, IM, as per Baker et al. 1990). Length and girth were measured when handling seals for tagging and used to post-calculate the weight (Castellini and Kooyman, 1990) and better estimate the true dosage applied. After chemical restraint, the fur on the seals head was checked to evaluate moulting stage, and when adequate, a tag was fixed to the fur on the skull area, either using epoxy adhesive or high viscosity instant glue (Loctite, USA). To our knowledge, this is the first study ever to successfully use instant glue for deploying tags on seals in Antarctica.

### Environmental covariates data

Environmental covariates values were obtained by spatially overlaying presences/pseudo-absence locations on spatial raster objects using the R software. For ocean depth, the IBCSO v2 dataset (Dorschel et al. 2022) was used. Monthly averaged ocean bottom temperature and surface-ice fraction were obtained from E.U. Copernicus Marine Service Information (downloaded on 1^st^ September 2022): a reanalysis dataset, Global Ocean Physical Multi Year product (doi.org/10.48670/moi-00021) was used for locations dated up to May 2020; a forecast dataset, GLOBAL Ocean Sea Physical Analysis and Forecasting Products (doi.org/10.48670/moi-00016), was used for the remaining locations, dated form May 2020 to July 2022. The distance from locations to the coastline was measured using a map available from the UK Polar Data Centre, Natural Environment Research Council of the UK (Gerrish, L., Fretwell, P., & Cooper 2020).

### Tracking data pre-processing – data filtering

Seal tracking data, obtained via the Argos system (CLS, Collective Localization Satellites, France/U.S.), were first filtered to retain only locations associated with speeds of 2 m/s or less. Because many of the Argos locations had high uncertainty (Vincent et al. 2002), and because consecutive locations were not separated regular time intervals, R software package *foieGras* (version 0.7; Jonsen et al. 2019) was used to re-estimate seal tracks in 12-hour intervals. After that, to avoid spatial bias from concentration of locations around tagging locations, only tracks longer than 20 days were retained for modelling. One male Weddell seal tagged in 2020, “Pendragon”, was the only animal to go beyond the continental shelf (Supplementary Fig. 2). For that reason, that track was removed from the analysis to not influence the spatial range of accessibility polygons (see *Modelling*) in a way that would not represent the background environment accessible for the remaining seals.

### Modelling

To represent the available background habitat in a presence/pseudo-absence approach to investigate Weddell seals habitat use, five random pseudo-absences per presence were created within accessibility polygons, i.e., minimum convex polygons created around all post-processed seal locations (Aarts et al. 2008), with a 0.2 arc-degree buffer. To minimize biased representation of environmental covariate values, due to covariate gaps within the spatiotemporal range of the presence/pseudo-absence dataset, whenever a presence or its corresponding five pseudo-absences had no covariate values, that entire group of six points was removed from the dataset. Anticipating the possibility of many pseudo-absences to be placed in areas with no coverage for “ocean bottom temperature” (identified to be important in preliminary modelling), accessibility polygons were edited to exclude regions with no values for that, before creating pseudo-absences. Since seal tracks in 2019 and 2020 did not spatially overlap with those in 2022, two accessibility polygons were created (Supplementary Figure 1). The sensitivity of models to the presence-pseudo-absence ratio was tested by fitting models with alternative ratios (i.e., 1/1 and 1/3), which indicated 1/5 to be adequate.

Factor covariates tested were month (9 levels: February to October), year (3 levels: 2019, 2020 and 2022) and the accessibility polygon (2 levels: polygon 1, with location from 2019 and 2020; polygon 2, with locations from 2022). Because the accessibility polygon was dictated by the sampling year, those two covariates were tested in separate candidate models. No highly correlated covariates (Supplementary Fig. 3) were tested simultaneously in models.

Binomial models with logit link function were fitted in a modelling framework combining Generalised Estimating Equations (GEEs; Hardin and Hilbe 2002) and Generalised Additive Models (GAMs; Wood, 2017), using software R (R core team, 2020), similar to the approach employed on Pirotta et al. (2011). The data were organised in correlation panels, with a panel for each set of presences within a track (one panel per track) and a different panel for each pseudo-absence (one panel per pseudo-absence). Using this panel structure was congruent with the assumption that locations within a track were correlated, but that locations in different tracks were not, and that pseudo-absences were mutually independent. To account for the imbalance between the number of presences and pseudo-absences, pseudo-absences were given 1/5th the weight of presences.

SALSA (Spatially Adaptive Local Smoothing Algorithm; Walker et al. 2011) was implemented with the MRSea R package (version 1.3.1; Scott-Hayward et al. 2013) to automatically select the number and locations of knots for smooth (b-splines) functions of numeric covariates in the models. The maximum number of knots were restricted to six to prevent overfitting of smooth terms (Wood, 2017). Covariate selection was conducted, first by removing smoothed covariates for which no knot was indicated as statistically significant (at α = 0.05), according to the robust standard errors (i.e., default output of MRSea package). Secondly, models were fitted as GEEs (geepack R package, version 1.3-2; (Halekoh, Højsgaard, and Yan 2006) to accommodate residual autocorrelation. At this stage, a backward step covariate selection was conducted: starting with the full model (i.e., the one with all covariates retained after the first step of covariate selection), a set of models was fitted, each with one less covariate, and models were compared using the QICu score (Pan 2001) to select the model to start the next covariate selection step. That was repeated until that score could not be improved. The significance of covariates retained in the final model was verified using function “getPvalues” from MRSea R package (Scott-Hayward et al. 2013).

Model performance was verified with Receiver Operating Characteristic (ROC) curves and confusion matrices, using the R package ROCR (version 1.0-7; Sing et al. 2005). The ROC and confusion matrix were used to calculate percentages of false positives and false negatives expected for the model, by comparing the predicted values to the observed. The contribution of each covariate in the final model was visualized with partial plots, with confidence intervals based on the GEE estimated uncertainty (Pirotta et al. 2011). For mapping Weddell seal distribution, the selected model was used to predict the probability of animal occurrence over a grid of locations spaced 5 km apart within the accessibility polygons. The time-varying covariate values for ice fraction and temperature used in predictions were extracted from the same datasets as for those in the model and grouped by months using the average of monthly observations in the last 10 years (e.g., 2012 to 2022).

## Author contribution

The study was conceptualized by GAB and LB. Data collection was performed by LB and GB. Methodology, data processing and analysis were done by GAB. GAB wrote the original draft. LB and KJH were responsible for funding acquisition GAB, LB and KJH revised and approved the final version of the manuscript.

## Acknowledgements

We thank all researchers and support staff in the NBP research cruises for assistance in data collection. We are especially grateful to Mark Barham, Tiago Dotto, Bastien Queste, Callum Rollo, Tasha Snow, Natalie Swaim, Hannah Wyles and Yixi Zheng for direct assistance with seal tagging, and to Robert Larter, Julia Wellner, Patricia Yager and Rob Hall for leading the research cruises. This work is part of the TARSAN project, a component of the International Thwaites Glacier Collaboration (ITGC Contribution No. ITGC-085), and supported by the National Science Foundation (NSF: Grant 1929991) and the Natural Environment Research Council (NERC: Grant NE/S006419/1 and NE/S006591/1). Fieldwork logistics were provided by NSF-U.S. Antarctic Program and NERC-British Antarctic Survey.

## References

Aarts, Geert, Monique MacKenzie, Bernie McConnell, Mike Fedak, and Jason Matthiopoulos. 2008. “Estimating Space-Use and Habitat Preference from Wildlife Telemetry Data.” Ecography 31 (1): 140–60. https://doi.org/10.1111/j.2007.0906-7590.05236.x.

Ainley, David G., Paul A. Cziko, Nadav Nur, Jay J. Rotella, Joseph T. Eastman, Michelle Larue, Ian Stirling, and Peter A. Abrams. 2021. “Further Evidence That Antarctic Toothfish Are Important to Weddell Seals.” Antarctic Science 33 (1): 17–29. https://doi.org/10.1017/S0954102020000437.

Baker, J. R., M. A. Fedak, S. S. Anderson, T. Arnborm, and R. Baker. 1990. “Use of a Tiletamine-Zolazepam Mixture to Immobilise Wild Grey Seals and Southern Elephant Seals.” Veterinary Record 126 (4): 75–77. https://doi.org/10.1136/vr.126.4.75.

Bengtson, John L., Jeff L. Laake, Peter L. Boveng, Michael F. Cameron, M. Bradley Hanson, and Brent S. Stewart. 2011. “Distribution, Density, and Abundance of Pack-Ice Seals in the Amundsen and Ross Seas, Antarctica.” Deep-Sea Research Part II: Topical Studies in Oceanography 58 (9–10): 1261–76. https://doi.org/10.1016/j.dsr2.2010.10.037.

Boehme, L., P. Lovell, M. Biuw, F. Roquet, J. Nicholson, S. E. Thorpe, M. P. Meredith, and M. Fedak. 2009. “Animal-Borne CTD-Satellite Relay Data Loggers for Real-Time Oceanographic Data Collection.” Ocean Science Discussions 6 (2): 1261–87. https://doi.org/10.5194/osd-6-1261-2009.

Börger, Luca, Novella Franconi, Giampiero De Michele, Alberto Gantz, Fiora Meschi, Andrea Manica, Sandro Lovari, and Tim Coulson. 2006. “Effects of Sampling Regime on the Mean and Variance of Home Range Size Estimates.” Journal of Animal Ecology 75 (6): 1393–1405. https://doi.org/10.1111/j.1365-2656.2006.01164.x.

Castellini, Michael A., and Gerald L. Kooyman. 1990 “Length, girth and mass relationships in Weddell seals (Leptonychotes weddellii).” Marine Mammal Science 6: 75–77.

CCAMRL. 2022. “Convention Area.” 2022. https://www.ccamlr.org/en/organisation/convention-area.

Cozannet, Gonéri Le, Thomas Bulteau, Bruno Castelle, Roshanka Ranasinghe, Guy Wöppelmann, Jeremy Rohmer, Nicolas Bernon, Déborah Idier, Jessie Louisor, and David Salas-y-Mélia. 2019. “Quantifying Uncertainties of Sandy Shoreline Change Projections as Sea Level Rises.” Scientific Reports 9 (1): 1–11. https://doi.org/10.1038/s41598-018-37017-4.

DeConto, Robert M., and David Pollard. 2016. “Contribution of Antarctica to Past and Future Sea-Level Rise.” Nature 531 (7596): 591–97. https://doi.org/10.1038/nature17145.

Dorschel, Boris, Laura Hehemann, Sacha Viquerat, Fynn Warnke, Simon Dreutter, Yvonne Schulze Tenberge, Daniela Accettella, et al. 2022. “The International Bathymetric Chart of the Southern Ocean Version 2.” Scientific Data 9 (1): 1–13. https://doi.org/10.1038/s41597-022-01366-7.

Dotto, Tiago Segabinazzi, Karen Heywood, Rob Hall, Tasha Snow, Christian Wild, and Lars Boehme. n.d. “Ocean Variability beneath Thwaites Eastern Ice Shelf Driven by the Pine Island Bay Gyre Strength.”

Gerrish, L., Fretwell, P., & Cooper, P. 2020. “High Resolution Vector Polygons of the Antarctic Coastline (7.3) [Data Set].” https://doi.org/https://doi.org/10.5285/ee7c6af2-da57-4519-8637-812eec5ff782.

Goetz, Kimberly T., Jennifer M. Burns, Luis A. Hückstӓdt, Michelle R. Shero, and Daniel P. Costa. 2017. “Temporal Variation in Isotopic Composition and Diet of Weddell Seals in the Western Ross Sea.” Deep-Sea Research Part II: Topical Studies in Oceanography 140: 36–44. https://doi.org/10.1016/j.dsr2.2016.05.017.

Halekoh, Ulrich, Søren Højsgaard, and Jun Yan. 2006. “The R Package Geepack for Generalized Estimating Equations.” Journal of Statistical Software 15 (2): 1–11. https://doi.org/10.18637/jss.v015.i02.

Hanchet, Stuart, Keith Sainsbury, Doug Butterworth, Chris Darby, Viacheslav Bizikov, Olav Rune Godø, Taro Ichii, Karl Hermann Kock, Luis López Abellán, and Marino Vacchi. 2014. “CCAMLR’s Precautionary Approach to Management Focusing on Ross Sea Toothfish Fishery.” Antarctic Science 27 (4): 333–40. https://doi.org/10.1017/S095410201400087X.

Hardin, James W., and Joseph M. Hilbe. 2002. Generalized Estimating Equations. Chapman and Hall/CRC. https://doi.org/10.1201/9781420035285.

Hogan, Kelly, Robert Larter, Alastair Graham, Robert Arthern, James Kirkham, Rebecca Totten Minzoni, Tom Jordan, et al. 2020. “Revealing the Former Bed of Thwaites Glacier Using Sea-Floor Bathymetry.” The Cryosphere Discussions, 1–36. https://doi.org/10.5194/tc-2020-25.

Jonsen, I. D., C. R. McMahon, T. A. Patterson, M. Auger-Méthé, R. Harcourt, M. A. Hindell, and S. Bestley. 2019. “Movement Responses to Environment: Fast Inference of Variation among Southern Elephant Seals with a Mixed Effects Model.” Ecology 100 (1). https://doi.org/10.1002/ecy.2566.

LaRue, Michelle, Leo Salas, Nadav Nur, David Ainley, Sharon Stammerjohn, Jean Pennycook, Melissa Dozier, et al. 2021. “Insights from the First Global Population Estimate of Weddell Seals in Antarctica.” Science Advances 7 (39). https://doi.org/10.1126/sciadv.abh3674.

McMahon, C. R., H. Burton, S. Mclean, D. Slip, and M. Bester. 2000. “Field Immobilisation of Southern Elephant Seals with Intravenous Tiletamine and Zolazepam.” Veterinary Record 146 (9): 251–54. https://doi.org/10.1136/vr.146.9.251.

Pan, Wei. 2001. “Akaike’s Information Criterion in Generalized Estimating Equations.” Biometrics 57 (1): 120–25. https://doi.org/10.1111/j.0006-341X.2001.00120.x.

Paolo, Fernando S., Helen A. Fricker, and Laurie Padman. 2015. “Volume Loss from Antarctic Ice Shelves Is Accelerating.” Science 348 (6232): 327–31. https://doi.org/10.1126/science.aaa0940.

Pirotta, Enrico, Jason Matthiopoulos, Monique MacKenzie, Lindesay Scott-Hayward, and Luke Rendell. 2011. “Modelling Sperm Whale Habitat Preference: A Novel Approach Combining Transect and Follow Data.” Marine Ecology Progress Series 436: 257–72. https://doi.org/10.3354/meps09236.

Queirós, José P., Darren W. Stevens, Matthew H. Pinkerton, Rui Rosa, Bernardo Duarte, Alexandra Baeta, Jaime A. Ramos, and José C. Xavier. 2022. “Feeding and Trophic Ecology of Antarctic Toothfish Dissostichus Mawsoni in the Amundsen and Dumont D’Urville Seas (Antarctica).” Hydrobiologia 849 (10): 2317–33. https://doi.org/10.1007/s10750-022-04871-3.

R Core Team. 2020. “R: A language and environment for statistical computing.” R Foundation for Statistical Computing, Vienna, Austria. URL https://www.R-project.org/.

Rignot, Eric, Jérémie Mouginot, Bernd Scheuchl, Michiel Van Den Broeke, Melchior J. Van Wessem, and Mathieu Morlighem. 2019. “Four Decades of Antarctic Ice Sheet Mass Balance from 1979–2017.” Proceedings of the National Academy of Sciences of the United States of America 116 (4): 1095–1103. https://doi.org/10.1073/pnas.1812883116.

Rotella, Jay J., William A. Link, Thierry Chambert, Glenn E. Stauffer, and Robert A. Garrott. 2012. “Evaluating the Demographic Buffering Hypothesis with Vital Rates Estimated for Weddell Seals from 30years of Mark-Recapture Data.” Journal of Animal Ecology 81 (1): 162–73. https://doi.org/10.1111/j.1365-2656.2011.01902.x.

Scott-Hayward, Lindesay, Cornelia Oedekoven, Monique Mackenzie, and Eric Rexstad. 2013. “Using the MRSea Package.”

Sing, Tobias, Oliver Sander, Niko Beerenwinkel, and Thomas Lengauer. 2005. “ROCR: Visualizing Classifier Performance in R.” Bioinformatics 21 (20): 3940–41. https://doi.org/10.1093/bioinformatics/bti623.

Vincent, Cécile, Bernie J. Mcconnell, Vincent Ridoux, and Michael A. Fedak. 2002. “Assessment of Argos Location Accuracy from Satellite Tags Deployed on Captive Gray Seals.” Marine Mammal Science 18 (1): 156–66. https://doi.org/10.1111/j.1748-7692.2002.tb01025.x.

Walker, C. G., M. L. Mackenzie, C. R. Donovan, and M. J. O’Sullivan. 2011. “SALSA – a Spatially Adaptive Local Smoothing Algorithm.” Journal of Statistical Computation and Simulation 81 (2): 179–91. https://doi.org/10.1080/00949650903229041.

Wheatley, Kathryn E., Corey J.A. Bradshaw, Robert G. Harcourt, Lloyd S. Davis, and Mark A. Hindell. 2006. “Chemical Immobilization of Adult Female Weddell Seals with Tiletamine and Zolazepam: Effects of Age, Condition and Stage of Lactation.” BMC Veterinary Research 2: 1–8. https://doi.org/10.1186/1746-6148-2-8.

Wood, Simon N. “Generalized additive models: an introduction with R”. chapman and hall/CRC, 2006.

Yoon, Seung Tae, Won Sang Lee, Sung Hyun Nam, Choon Ki Lee, Sukyoung Yun, Karen Heywood, Lars Boehme, et al. 2022. “Ice Front Retreat Reconfigures Meltwater-Driven Gyres Modulating Ocean Heat Delivery to an Antarctic Ice Shelf.” Nature Communications 13 (1): 1–8. https://doi.org/10.1038/s41467-022-27968-8.

Zheng, Yixi, Karen J. Heywood, Benjamin G.M. Webber, David P. Stevens, Louise C. Biddle, Lars Boehme, and Brice Loose. 2021. “Winter Seal-Based Observations Reveal Glacial Meltwater Surfacing in the Southeastern Amundsen Sea.” Communications Earth and Environment 2 (1): 1–9. https://doi.org/10.1038/s43247-021-00111-z.

