## Supplementary for "Weddell seals near the fastest melting glacier in Antarctica prefer shallow, coastal and partially ice-covered waters"

for

by

Guilherme A. Bortolotto, Karen J. Heywood, Lars Boehme

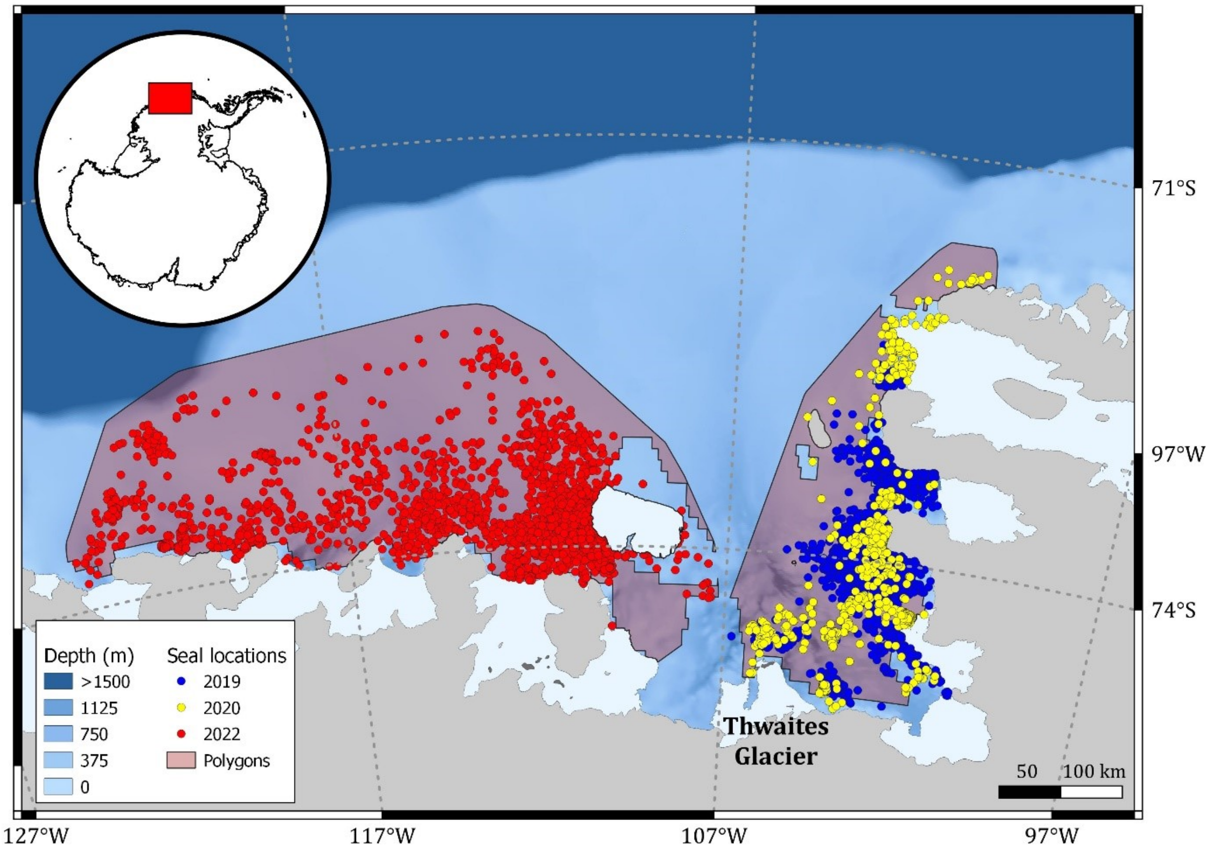


**Supplementary Fig. 1. Seal tracking data** – post-processed seal locations in 2019, 2020 and 2022 used to model habitat use of Weddell seals in the Amundsen Sea, West Antarctica. "Polygons" refers to the accessibility polygons within which pseudo-absences were randomly created.

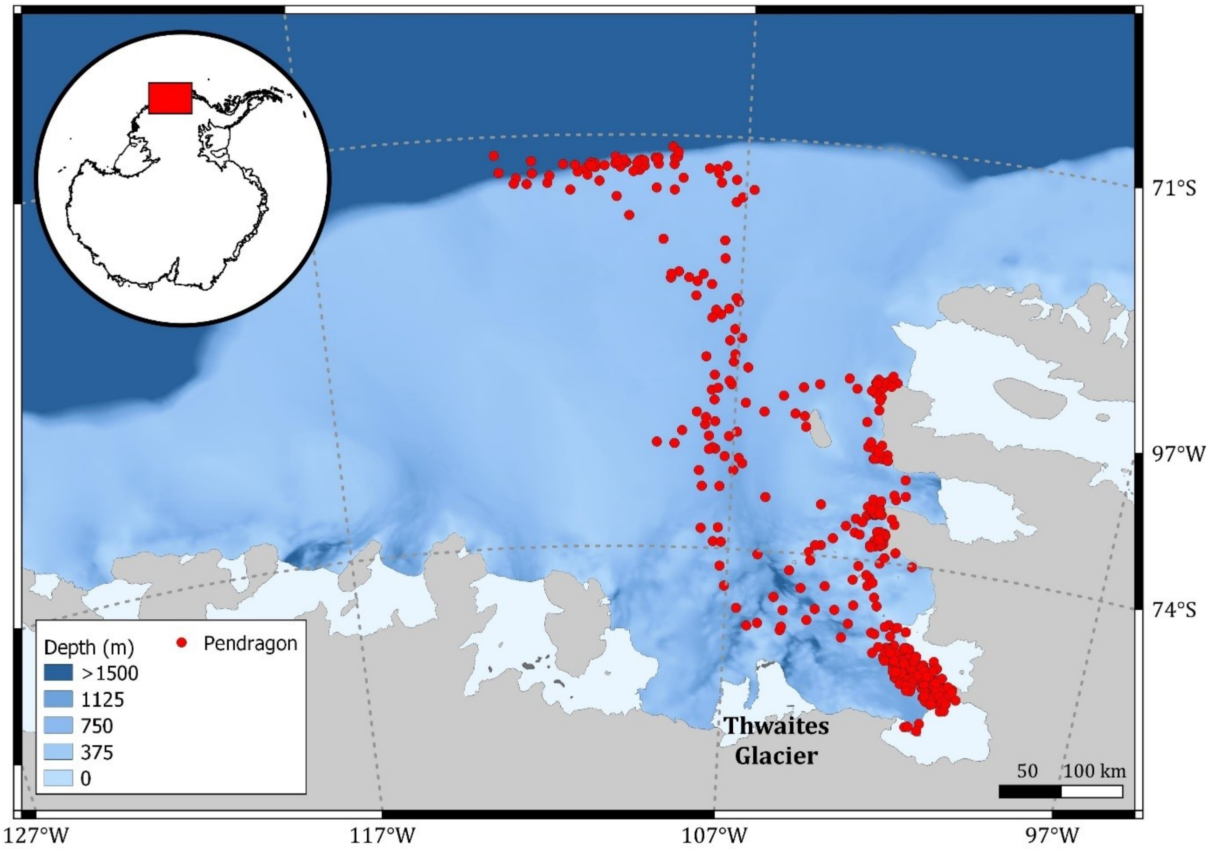


**Supplementary Fig. 2. Pendragon track** – post-processed locations for Pendragon, a seal tagged in 2020, removed from the distribution and habitat use analysis because it was the only seal in the dataset to go beyond the continental shelf. Its inclusion would have caused a large expansion of the accessibility polygon for the seal locations in 2019 and 2020, that would not represent the overall pattern.

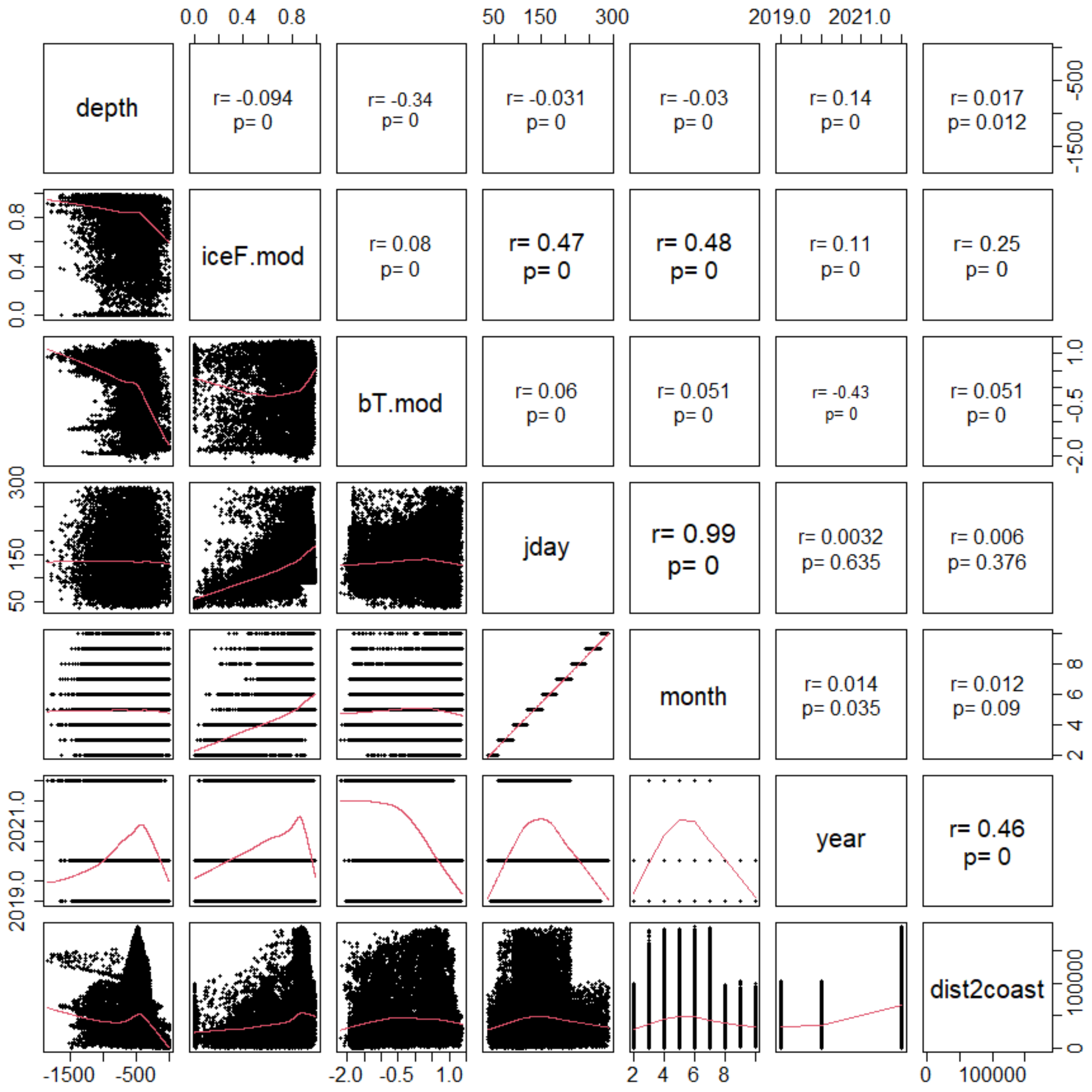


**Supplementary Fig. 3. Covariates correlation** – Linear correlation diagnostic plots for covariates considered for habitat use modelling of Weddell seals in the Amundsen Sea.

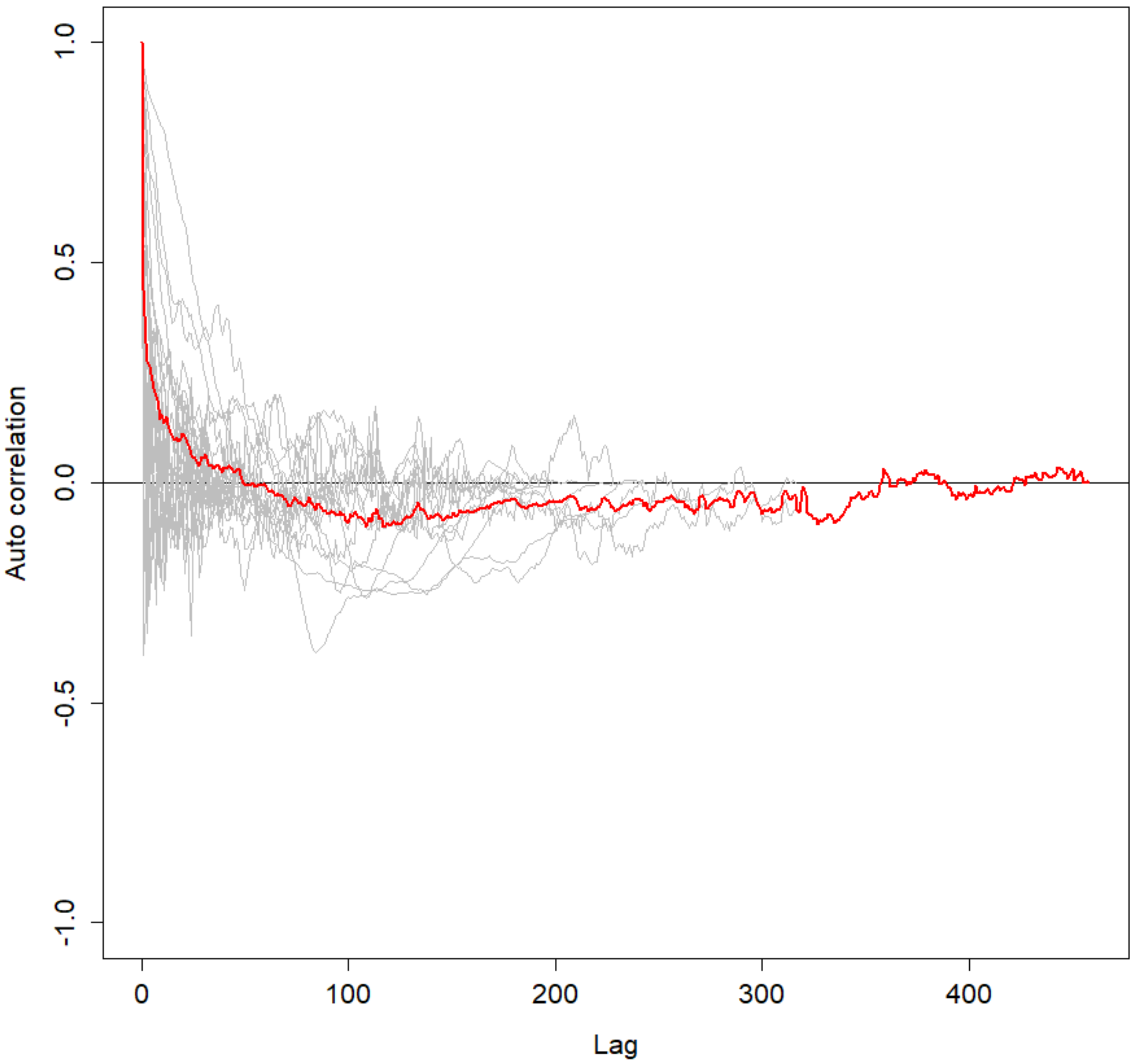


**Supplementary Fig. 4. Model residual auto-correlation** – The model fitted with SALSA in the first step of covariate selection, indicated very high residual autocorrelation, illustrated using function “runACF” from R package MRSea (version 1.3.1; Scott-Hayward et al., 2013)

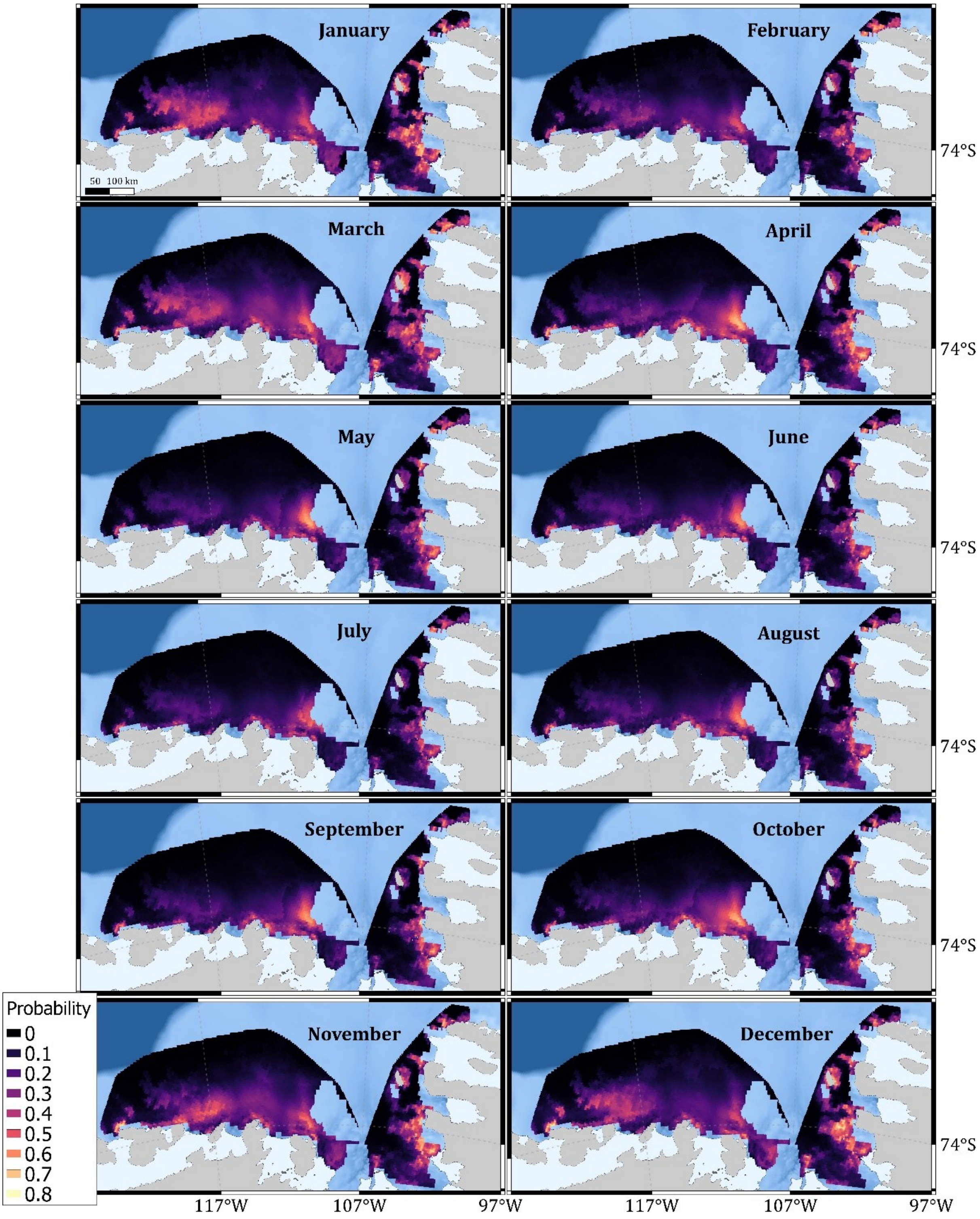


**Supplementary Fig. 5. Weddell seal distribution predictions for each month** – Note that seal locations used in the models did were not available for January and December, so maps for those months are extrapolations beyond the time range observed.

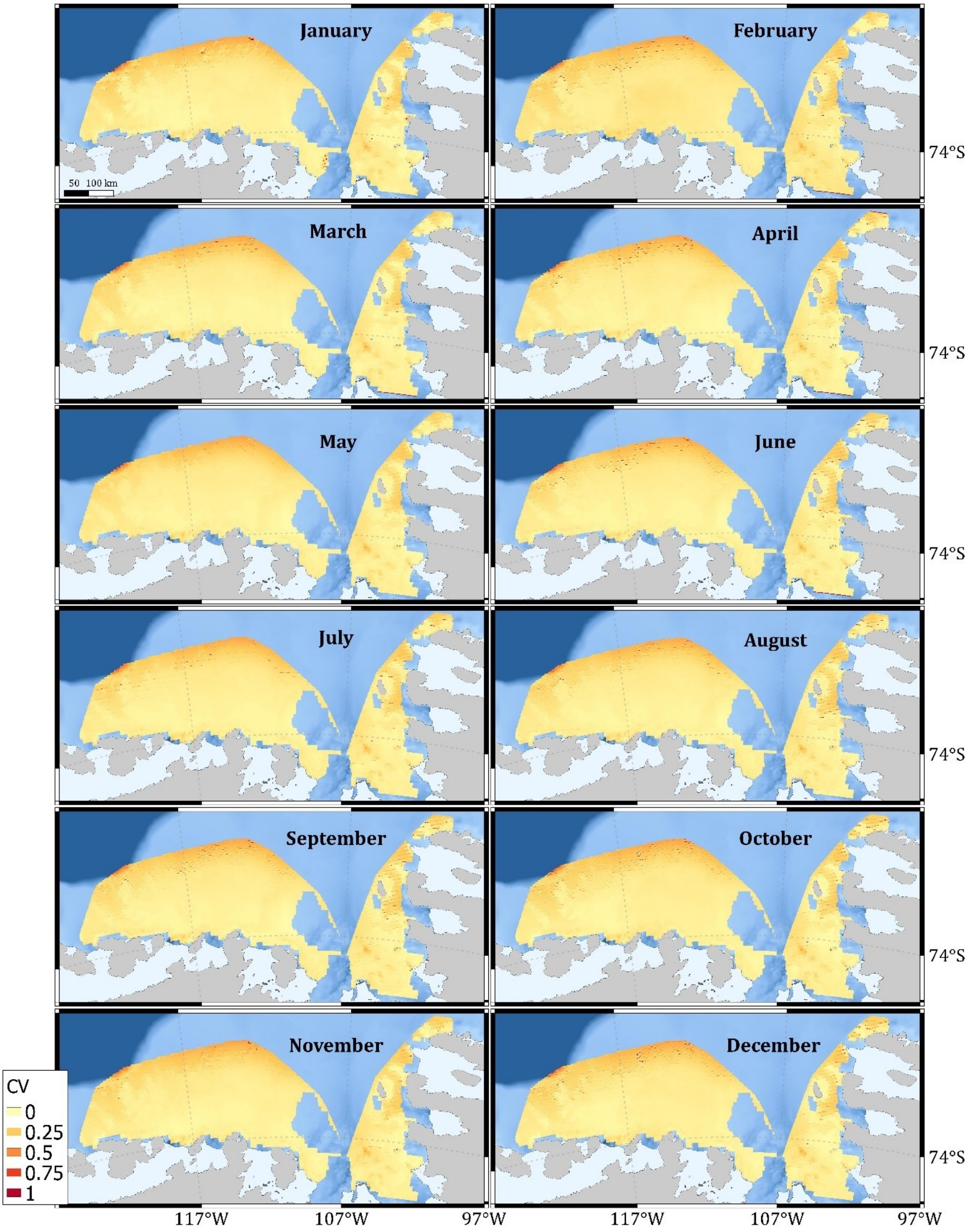


**Supplementary Fig. 6. Uncertainty in Weddell seal distribution predictions for each month** – Coefficients of variation (CV) were calculated as the ratio of seal occurrence probability standard errors and probability estimates.
